# pH-dependent spontaneous hydrolysis rather than gut bacterial metabolism reduces levels of the ADHD treatment, Methylphenidate

**DOI:** 10.1101/2020.07.06.189191

**Authors:** Julia Aresti-Sanz, Walid Maho, Rob Rodrigues Pereira, Hjalmar Permentier, Sahar El Aidy

## Abstract

Methylphenidate is absorbed in the small intestine. The drug is known to have low bioavailability and a high interindividual variability in terms of response to the treatment. Gut microbiota has been shown to reduce the bioavailability of a wide variety of orally administered drugs. Here, we tested the ability of small intestinal bacteria to metabolize methylphenidate. *In silico* analysis identified several small intestinal bacteria to harbor homologues of the human carboxylesterase 1 enzyme responsible for the hydrolysis of methylphenidate in the liver. Despite our initial results hinting towards possible bacterial hydrolysis of the drug, up to 60% of methylphenidate was spontaneously hydrolyzed in the absence of bacteria and this hydrolysis was pH-dependent. Overall, the study shows that pH-dependent spontaneous hydrolysis rather than gut bacterial metabolism reduces levels of methylphenidate and suggest a role of the luminal pH in the bioavailability of the drug.

## Introduction

Attention-deficit/hyperactivity disorder (ADHD) is one of the most prevalent neurodevelopmental disorders, affecting 6-12% of children and persisting into adulthood in around 60% of the cases [1]. Although a cause-effect relationship has not yet been established for ADHD, altered levels of dopamine and norepinephrine, and their corresponding transporters in the brain, seem to play a key role in the cognitive impairment and dysregulated reward system that characterize ADHD [2,3]. Thus, ADHD is mainly treated with amphetamine-like psychostimulants that improve symptoms by increasing the levels of dopamine and norepinephrine neurotransmitters in the brain.

Methylphenidate (MPH), a dopamine reuptake inhibitor, is considered as the golden treatment for ADHD [4,5]. MPH is administered orally and rapidly absorbed into the blood steam through the small intestine (SI), reaching peak concentrations between 1- and 3-h post ingestion. Around 70% of MPH is recovered in urine in the form of ritalinic acid (RA). RA is the inactive metabolite of MPH produced in the liver by carboxylesterase 1 (CES1) [6]. Despite its efficacy, MPH has a low bioavailability of around 30%. Moreover, there is a high interindividual variability among patients in terms of their response to the treatment [7,8].

First-pass metabolism could explain the low bioavailability of MPH. Genetic variations in *CES1* have been shown to impact enzyme activity towards different substrates *in vitro* [9], which could account for differences in MPH hydrolysis among patients. Nevertheless, clinical human studies are scarce. Increased concentrations of MPH were found in plasma of individuals carrying a polymorphism in the *CES1* gene indicating decreased enzyme activity [10,11]. However, this study was performed on a small number of healthy volunteers administered a single dose of MPH, which cannot be translated to ADHD patients on multiple doses per day of the drug. Recently, absorption of the drug was modelled based on physicochemical properties of the drug, formulation-related information, and differences in gut physiology along the gastrointestinal tract [12]. Importantly, non-specific intestinal loss of MPH had to be introduced in the model in order to obtain plasma profiles of MPH and RA comparable to those found in clinical studies. Thus, the model suggested intestinal loss of MPH prior to absorption and hepatic/systemic metabolism [12][13]. Nonetheless, the mechanism explaining such non-specific intestinal loss remains to be explained, since the CES1 enzyme is absent in the gastrointestinal tract [14].

The gut microbiota represents a metabolic factory able to synthesize indigenous and exogenous compounds, such as food components and drugs, in the host [15][16][17][18]. Bacterial metabolism of MPH could therefore explain the potential intestinal loss of MPH. Indeed, bacterial esterases that are able to hydrolyze carboxyl esters have been previously described [18]. For example, the highly abundant gut bacterium *Escherichia coli* harbors the esterase yjfP, which is able to hydrolyze the ester 4-nitrophenylacetate [19]. Similarly, *Bacillus subtilis* pnbA esterase has been shown to hydrolyze 4-nitrophenylacetate [20]. The present study investigates whether gut bacteria can metabolize MPH leading to increased presystemic metabolism and reduced bioavailability of the drug, thereby interfering with the efficacy of MPH medication in ADHD patients.

## Material and methods

### Rat luminal content

Luminal small intestinal content of wild-type Groningen (WTG) rats (n = 5) was collected in sterile Eppendorf tubes by gentle pressing along the entire cecum and was snap frozen in liquid N_2_ and stored at −80 °C. 10% (w/v) suspensions of the luminal content were grown in enriched beef broth based on SHIME medium (**S1 Table)** [21]. Bacterial cultures within the inoculum were allowed to grow for 3 h, followed by supplementation with 50 μM Methylphenidate (MPH) hydrochloride tablets (10 mg, Mylan; provided by Dr. R. Pereira; Medical Center Kinderplein, Rotterdam, The Netherlands). MPH and were incubated at 37 °C in aerobic conditions with shaking at 220 rpm. Samples were collected at 0 and 24 h for HPLC-MS/MS analysis.

### Pure bacterial cultures

*L. salivarius* W1, *L. plantarum* W24 and *E. faecium* W54 strains were obtained from Winclove Probiotic B.V. *C. ammoniagenes* DSM20306, *E. coli* DSM1058 and *E. coli* DSM12250 were obtained from the German Collection of Microorganisms and Cell Cultures (DSMZ). Additionally, the lab strains *E. coli* BW25113 [22] and the vancomycin-resistant strain *E. faecalis* V583 [23] were used in this study. All bacterial strains were grown in incubators (New Brunswick Scientific) at 37 °C aerobically in in enriched beef broth (**S1 Table**). For experiments where the effect of pH on MPH hydrolysis was studied (**Fig 3**), culture media were prepared at different pH values. To do so, buffer solutions of KH_2_PO_4_/K_2_HPO_4_ were prepared at different concentrations to obtain the desired pH when adding them to the media. Strains which required shaking for proper growth (all *E. coli* strains, *C. ammoniagenes* DSM20306, *L. salivarius* W24 and *L. plantarum* W1) were grown with continuous agitation at 220 rpm. Bacteria were inoculated from −80 °C glycerol stocks and grown overnight. Before the experiment, cultures were diluted to 1% in fresh enriched beef broth medium and were grown until late exponential phase (**S2 Figure**). Growth was followed by measuring optical density (OD) at 600 nm in a spectrophotometer. 50 μM MPH was added to the cultures and samples were taken at 0 and 24 h for HPLC-MS/MS analysis.

**Fig 1.**
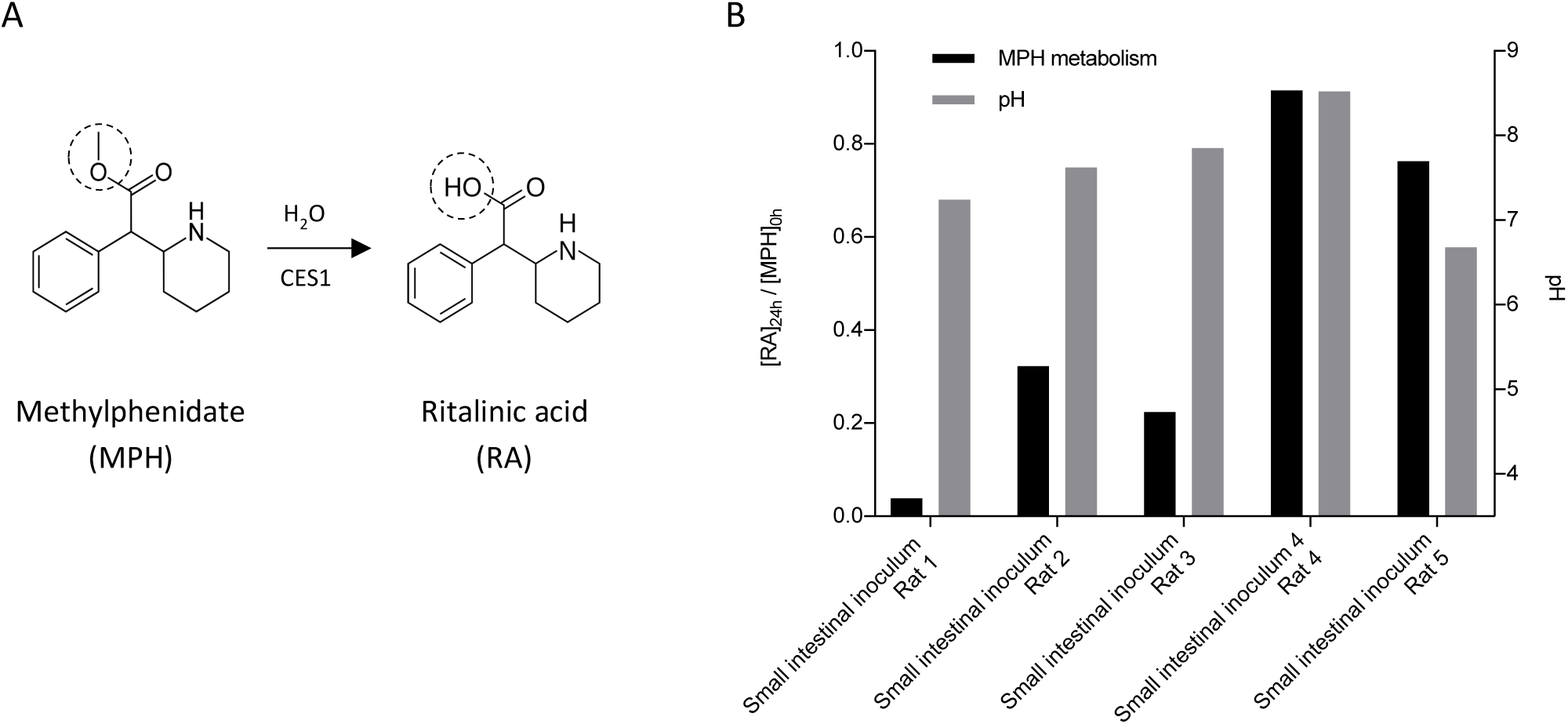
Methylphenidate (MPH) metabolism by small intestinal bacteria. **(A)** Hydrolysis reaction by CES, which removes a methyl group from MPH to form ritalinic acid (RA). **(B)** MPH metabolism by small intestinal luminal microbiota from WTG rats (n = 5) (black bars; left y-axis), and pH values measured in the cultures after 24 h of incubation with 50 μM MPH (grey bars; right y-axis). Metabolic activity is shown as the ratio of [RA]_24h_/[MPH]_0h_ quantified in ng/μL and normalized to d10-RA.

**Fig 2.**
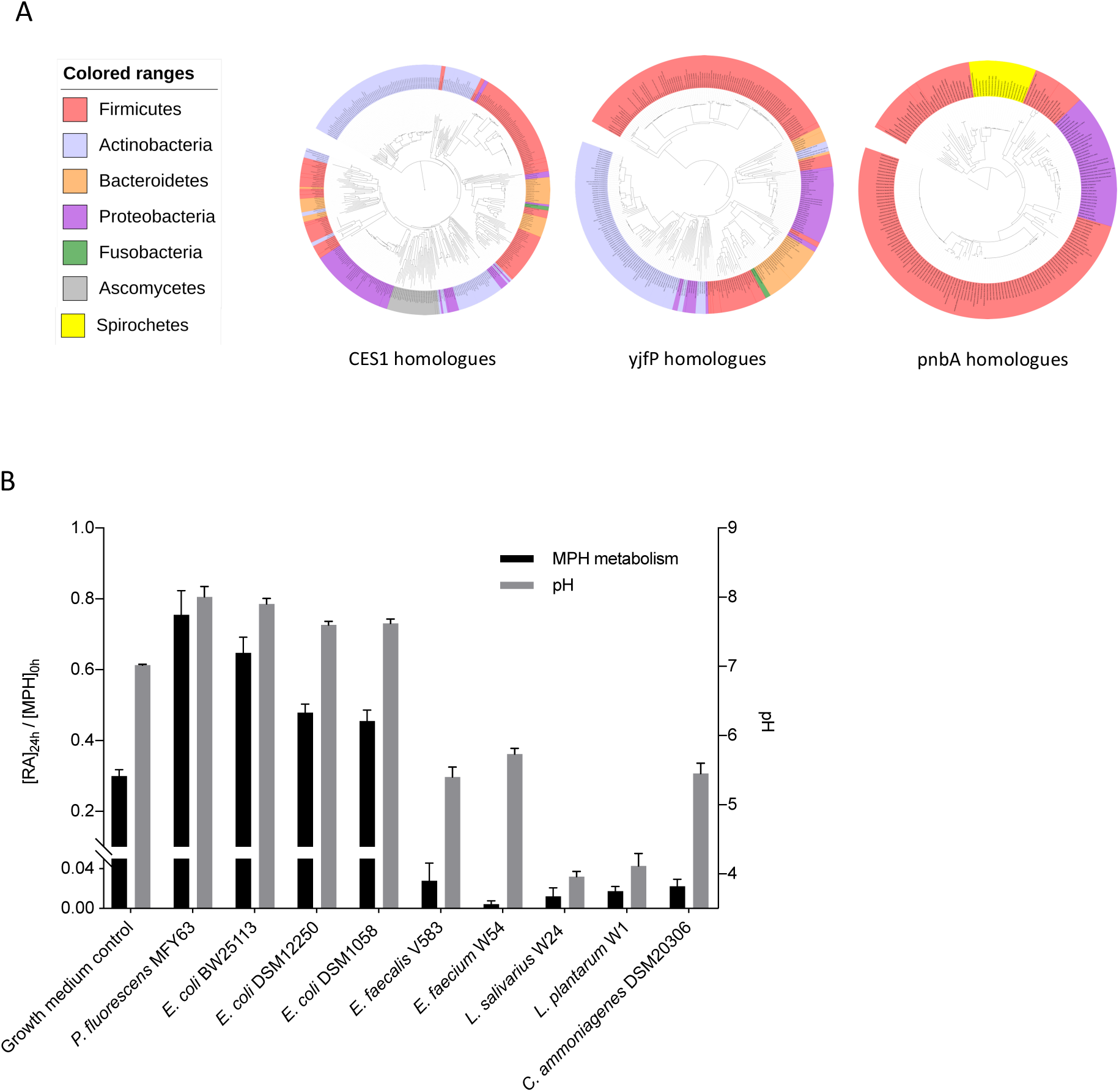
Methylphenidate (MPH) metabolism by pure bacterial cultures. **(A)** Phylogenetic trees created using iTOL online tool showing gut bacterial strains harboring homologue enzymes of human CES1, *E. coli* yjfP and *B. subtilis* pnbA respectively. **(B)** Screening of gut bacterial pure cultures for the metabolism of MPH. Metabolic activity is shown as the ratio of [RA]_24h_/[MPH]_0h_ quantified in ng/μL and normalized to d10-RA (black bars; left y-axis) together with pH measurements in the cultures after 24 h of incubation (grey bars; right y-axis) with 50 μM MPH. Error bars represent standard deviation (n = 3).

**Fig 3.**
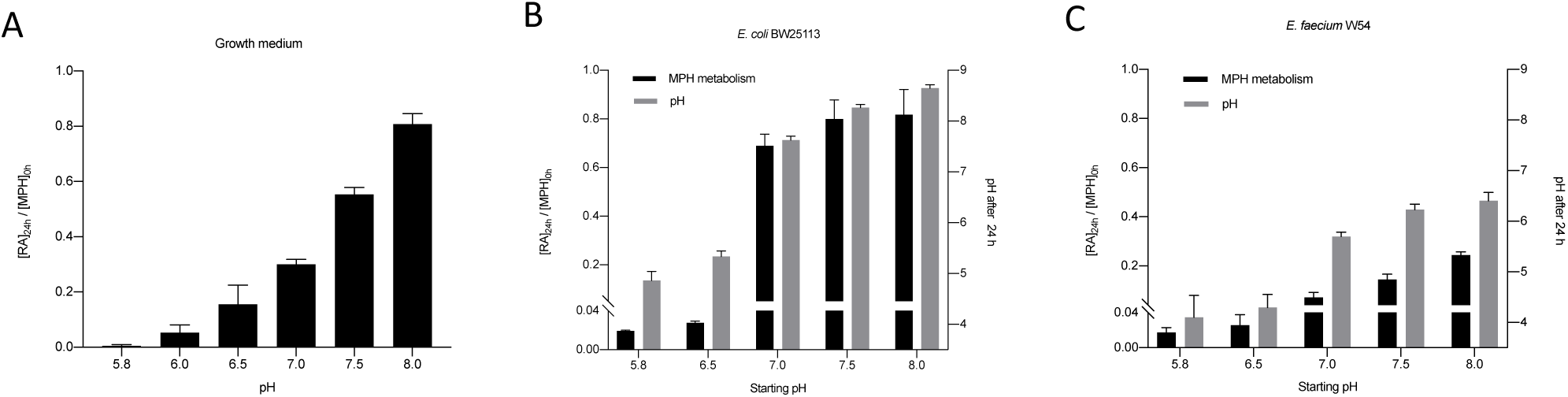
Effect of pH on methylphenidate hydrolysis. **(A)** MPH hydrolysis in growth medium prepared at different pH values. **(B, C)** pH effect on MPH hydrolysis in *E. faecium* W54 **(B)**, and *E. coli* BW25113 **(C)** (black bars; right y-axis) shown as the ratio of [RA]_24h_/[MPH]_0h_ quantified in ng/μL and normalized to d10-RA and pH measurements after 24 h of incubation with 50 μM MPH (grey bars; right y axis).

### HPLC-MS/MS sample preparation and analysis

In order to monitor the levels of MPH and Ritalinic acid (RA) hydrochloride solution (1 mg/mL as a free base, Sigma-Aldrich, The Netherlands) in bacterial cultures, samples were collected by adding 100 μL of culture to 400 μL of 100% methanol. The internal standard d10-Ritalinic acid (d10-RA) hydrochloride solution (100 μg/mL as a free base; Sigma-Aldrich, The Netherlands) was added to all samples at a final concentration of 2 ng/μL as an internal standard for accurate quantification. Samples were then centrifuged at 14000 rpm for 15 min at 4 °C. Supernatants were transferred to a clean tube and methanol was evaporated using a Savant speed-vacuum dryer (SPD131, Fisher Scientific, Landsmeer, Netherlands). Finally, samples were reconstituted in 500 μL of water.

Sample analysis was performed using a Shimazu HPLC system consisting of a SIL-20AC autosampler, a CTO-20AC column oven and LC-20AD liquid chromatograph pumps. Chromatography separation was achieved using a Waters CORTECS C18+ column (100×2.1 mm; 2.7 µm). The mobile phase consisted of a mixture of water (A) and acetonitrile (B) both containing 0.1% formic acid. A flow rate of 0.25 mL/min was used with a linear gradient: 5% (B) for 5 min, followed by an increase to 80% (B) in 5 min, which was kept for 3 min to wash the column and then returned to initial conditions for 2 min. The HPLC was coupled to an API3000 triple-quadrupole mass spectrometer (Applied Biosystems/MDS Sciex) via a turbo ion spray ionization source. Ionization was performed by electrospray in positive mode and selected reaction monitoring (SRM) was used to detect the metabolites. The SRM transitions were: m/z 234 to 84 for MPH, 220 to 84 for RA and 230 to 84 for d10-RA. Other parameters were set as follows for all transitions: declustering potential 15 V, entrance potential 7 V, focusing potential 65 V, collision energy 30 V and collision cell exit potential 14 V.

### Calibration standards and biological matrices

MPH standard was obtained by extraction from MPH hydrochloride tablets (10 mg) as follows; one tablet was crushed in a mortar and the resulting powder was diluted in 10 mL of a mixture containing acetonitrile, methanol, and acetate buffer pH 4 (0.2 M CH_3_COONa) in a ratio of 30:50:20, respectively. The solution was mixed with a magnetic stirrer for 10 min, and was allowed to stand until the solid phase containing the insoluble components of the tablets had precipitated. Next, the polar liquid phase containing MPH was collected with a syringe and sterilized using 0.2 µm filters. This resulted in stock solutions of 1 mg/mL of MPH which were stored at −20 °C until further use.

For MS quantification, calibration curves were obtained in different matrices to account for matrix effects in the detection of MPH and RA. Calibration samples containing MPH and RA in a concentration range of 0.01 to 5 ng/μL were prepared in methanol and 2 ng/μL of d10-RA was added as an internal standard to correct for intrasample variation of MPH and RA. Next, methanol was removed by vacuum centrifugation and samples were reconstituted in 500 μL of the relevant biological matrix. Two types of biological matrices were prepared. For quantification in pure bacterial cultures, *E. coli* BW25113 cultures were grown to late exponential phase (**S2 Figure**), cells were removed by centrifugation at 14000 rpm for 15 min and the supernatants were filtered and used for reconstitution. Similarly, a pool was made combining small intestinal content of 5 rats used in **Fig 1** to obtain a complex matrix for this experiment. The pooled inoculum was allowed to grow for 3 hours and supernatant was obtained by centrifugation and filtering as explained before to be used for reconstitution of the calibration curves. Linearity of the detection of MPH and RA in both biological matrices is shown in **S1 Figure**.

### Bioinformatics

Protein sequences of human CES1 (NCBI accession: AAI_10339.1), *E. coli* yjfP (NCBI accession: ANK_04958.1) and *B. subtilis* pnbA (NCBI accession: KIX_83209.1) were BLASTed against the protein sequences from the NIH Human Microbiome Project data bank using search limits for Entrez Query “43021[BioProject]”. All BLASTp hits were converted to a distance tree using MEGA5. The tree was exported in Newick format and visualized in iTOL phylogenetic display tool (http://itol.embl.de/).

### Statistical analysis

All statistical tests and linear regression models were performed using GraphPad Prism 7.

## Results

### Gut bacteria convert methylphenidate into ritalinic acid

Between 65-75% of MPH is absorbed in the small intestine [12]. To determine whether small intestinal bacteria have the ability to metabolize and inactivate MPH by hydrolysis of the ester group (**Fig 1A**), small intestinal luminal samples from wild-type Groningen rats (n = 5) were incubated aerobically *in vitro* with 50 µM MPH and the concentrations of MPH and RA were monitored by High-Performance Liquid Chromatography coupled with Tandem Mass Spectrometry (HPLC-MS/MS). Analytical details for the quantification method of both analytes are provided in **S1 Figure**. The concentration of MPH employed was based on the estimation that well-absorbed drugs are present in the small intestine at concentrations ≥ 20 µM [24].

Interestingly, there was a wide variation among the tested luminal samples in their ability to convert MPH into RA, ranging from samples that metabolized MPH to RA almost completely (90% metabolism), to samples where MPH was not metabolized to RA at all after 24 h (**Fig 1B**). The results suggest a role of gut bacteria in the conversion of MPH into RA. Gut microbiota may interfere with the bioavailable levels of MPH, which is absorbed in the upper gastrointestinal tract when taken as an ADHD medication.

### Gut bacteria harbor homologues for the human CES1 enzyme responsible for metabolization of methylphenidate

The human enzyme responsible for the hydrolysis of MPH to RA in the liver is CES1 [25]. We hypothesized that gut bacteria harbor a homologue for the human CES1 enzyme. To verify our hypothesis, the protein sequence (XP_005255831.1) from the human CES1 enzyme was used as a query to search the US National Center of Health Human Microbiome Project (HMP) protein database. The analysis identified several bacterial genera; *Corynebacterium, Bifidobacterium, Bacteroides, Klebsiella, Citrobacter* and *Faecalibacterium* to harbor highly homologous proteins to the human CES1 annotated as esterases or carboxylesterases (**Fig 2A**). Next, we focused our search on two known bacterial esterases from *E. coli* and *B. subtilis*. The protein sequence (ANK_04958.1) from *E. coli* yjfP esterase was used as a query to search the HMP protein database. The analysis identified bacteria belonging mainly to the Firmicutes phylum, as well as Proteobacteria. Specifically, *Enterococcus faecalis, Enterococcus faecium, Lactobacillus plantarum* and *Klebsiella* strains we found to harbor yjfP homologous proteins. Similarly, *Enterococcus* strains, *Faecalibacterium, Corynebacterium, Klebsiella, Citrobacter, Prevotella, Bacteroides, Bifidobacterium* and *Pseudomonas* were identified to harbor *B. subtilis* pnbA (KIX_83209.1) homologous proteins (**Fig 2A**).

Based on the *in-silico* analysis, a comprehensive screening of gut-associated bacterial strains harboring esterase proteins was performed. Out of all the bacteria found to harbor CES1, yjfP and pnbA homologues, we focused on gut bacteria known to inhabit the small intestine, the major site of MPH absorption [12]. To this end, pure cultures of the Gram-negative bacteria *Pseudomonas fluorescens* MFY63, *Escherichia coli* DSM12250, *E. coli* DSM1058, and the laboratory strain *E. coli* BW25113 were incubated aerobically with 50 µM of MPH and were analyzed by HPLC-MS/MS (details of the quantification method in these cultures are shown in **S1 Figure**). *P. fluorescens* MFY63, and *E. coli* BW25113 cultures displayed a conversion of 70% of MPH into RA after 24 h of aerobic incubation (**Fig 2B**). In the case of the gut isolates *E. coli* DSM1058 and *E. coli* DSM12250, 50% of MPH was hydrolyzed. In contrast, the Gram-positive bacteria *E. faecalis* V583, *E. faecium* W54, *L. plantarum* W1, *L. salivarius* W24, and *C. ammoniagenes* DSM20306 cultures did not metabolize MPH (**Fig 2B**) suggesting that certain Gram-negative bacteria are involved in the metabolism of MPH. Surprisingly, around 20% MPH was spontaneously hydrolyzed in the growth medium in the absence of bacteria, even higher than in the Gram-positive bacteria cultures.

### pH of the bacterial growth media causes hydrolysis of methylphenidate in the gut

The spontaneous hydrolysis of MPH observed in the bacteria growth medium in the absence of bacteria (**Fig 2B**) led us to investigate the role of pH in MPH hydrolysis. In bacterial cultures where MPH was not metabolized, the pH measured after 24 h ranged from 4.0 - 5.5. In contrast, bacterial cultures that showed high levels of MPH hydrolysis had a pH between 7.5 - 8.0. Moreover, the *E. coli* BW25113 cultures had a slightly higher average pH of 7.9 compared to *E. coli* DSM1058 and *E. coli* DSM12250, where the average pH was 7.5 and this was accompanied by a smaller percentage of MPH hydrolysis, 70% versus 50% respectively (**Fig 2B**). Indeed, Pearson r correlation analyses showed a strong positive correlation (r = 0.89, r^2^ = 0.79, P value = 0.0006) between MPH-hydrolyzing bacterial cultures and pH of the growth media. These findings suggest that pH of the bacterial culture, and not bacterial metabolic activity, is responsible for the hydrolysis of MPH into its inactive form, RA.

To further determine whether gut bacterial metabolic activity plays any role in the observed hydrolysis of MPH, MPH stability was tested in an enriched beef broth based on SHIME medium [21], the medium used in all our incubation experiments (**S3 Table**). Enriched beef broth was prepared at different pH values, ranging from 5.5 to 8.0 resembling the pH values previously measured in the different bacterial cultures (**Fig 2B**), and was incubated aerobically with 50 µM MPH for 24 h and analyzed by HPLC-MS/MS. At pH ≤ 6, which resembles the pH measured in bacterial cultures that did not hydrolyze MPH, ≥ 80% of MPH remained intact, while 80% of the drug was hydrolyzed to RA at pH 8 (**Fig 3A**). Pearson r correlation analyses showed a strong positive correlation (r = 0.98, r^2^ = 0.96, P value = 0.0005) between the pH value of the medium and the amounts of hydrolyzed MPH.

To determine whether the strong correlation found between pH and MPH hydrolysis could explain the differences in MPH metabolism in the bacterial pure cultures, we selected *E. coli* BW25113 that showed 70% hydrolysis of MPH into RA, and *E. faecium* W54 that did not hydrolyze the drug. *E. coli* BW25113 and *E. faecium* W54 were grown in enriched beef broth at pH values ranging from 5.5 to 8.0, incubated aerobically with 50 µM MPH for 24 h and analyzed by HPLC-MS/MS. Changes in pH after 24 h of incubation were measured and compared to the initial pH values.

When *E. coli* BW25113 was grown in enriched beef broth at pH ≤ 6.5, a negligible amount of MPH was hydrolyzed to RA after 24 h of incubation with 50 μM MPH and the pH of the 24 h culture dropped to 5.0-5.5. In contrast, when *E. coli* BW25113 was grown in enriched beef broth at pH ≥ 7 (the same pH of the culture plotted in **Fig 2B**), the pH of the culture rose to 7.5-8.5 after 24 h of incubation with MPH and 70-90% of MPH was hydrolyzed to RA (**Fig 3B**). On the other hand, when *E. faecium* W54 was grown in enriched beef broth at pH ≤ 6.5 the pH of the culture dropped below 5 after 24 h of incubation with MPH and a negligible amount of MPH was hydrolyzed to RA. When *E. faecium* W54 was grown at pH ≥ 7, the pH of the cultures dropped to values between 6.5-5.5 and this was accompanied by only 20% MPH hydrolysis (**Fig 3C**). Pearson r correlation analyses showed a positive correlation between the pH value of the *E. coli* BW25113 cultures (r = 0.90, r^2^ = 0.81, P value = 0.03), *E. faecium* W54 (r = 0.93, r^2^ = 0.8701, P value = 0.02) and the amounts of hydrolyzed MPH. Taken together, these results support the hypothesis that the MPH hydrolysis by gut bacteria observed in **Fig 2B** is likely to be due to changes in pH over the duration of bacterial incubation with MPH.

We next ruled out the possibility that certain components of the bacterial growth media could be catalyzing the hydrolysis of MPH. Medium composition **(S3 Table)** changes during the course of bacterial growth and their metabolism, which in turn, could deplete potential hydrolysis catalyzing agents. *E. coli* BW25113 and *E. faecium* W54 cultures were grown to late exponential phase **(S2 Figure)** and the supernatants were collected, filtered, and incubated with 50 µM MPH. pH values of *E. faecium* W54 supernatants, which were around 5.5, were adjusted to 6.0, 7.0 and 7.5, respectively, to resemble the pH previously measured in the different bacterial cultures **(Fig 2B)**. Interestingly, incubation of MPH with *E. faecium* W54 supernatants at pH 6 resulted in 10% hydrolysis of MPH to RA, but levels of hydrolysis increased with increasing pH values; 20-30% at pH 7 and 60% at pH 7.5, respectively **(Fig 4A)**. When *E. coli* BW25113 supernatants, which had a pH around 7.0 were adjusted to 7.5 to resemble the pH after 24h of growth (**Fig 2B**), the MPH hydrolysis increased from 20-30% at pH 7.0 (**Fig 2B**) to 60% at pH 7.5 (**Fig 4B**). Collectively, our results indicate that the majority of the observed hydrolyzed MPH results from pH-dependent spontaneous non-enzymatic conversion rather than from bacterial metabolism.

**Fig 4.**
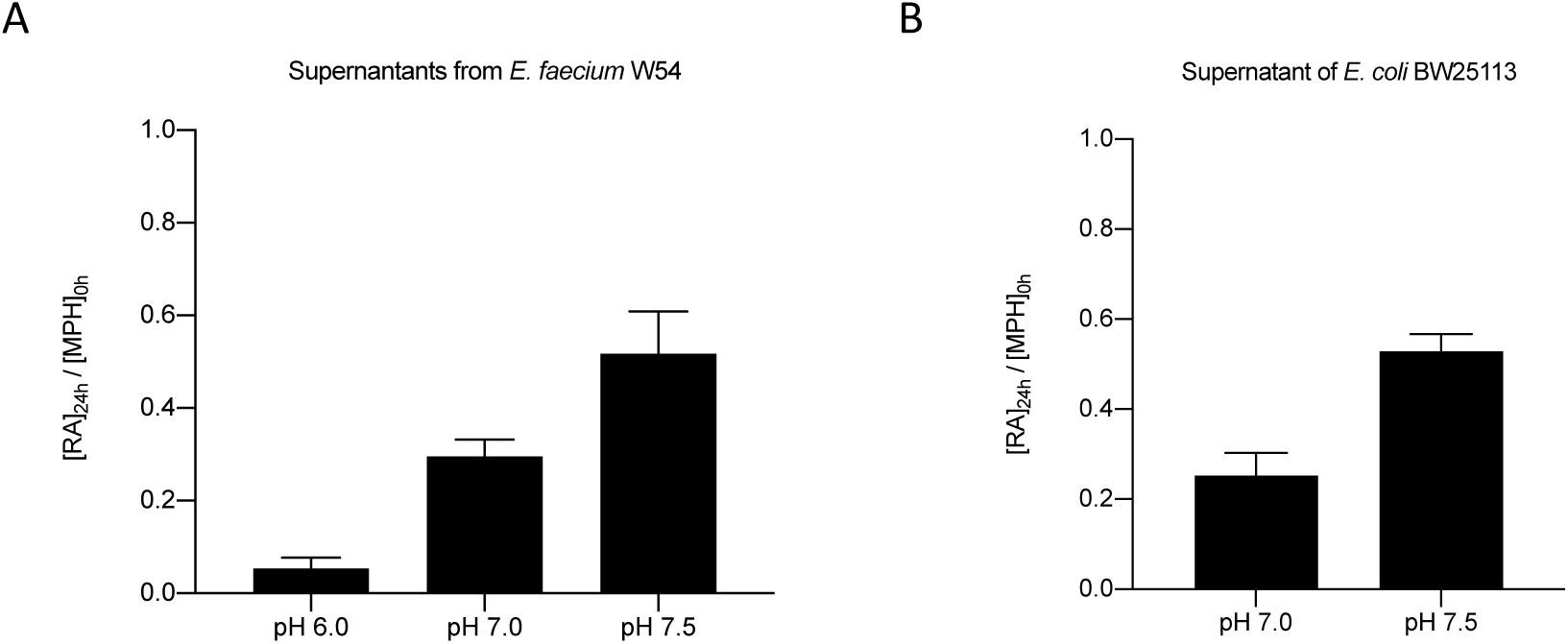
Methylphenidate hydrolysis in bacterial supernatants. MPH hydrolysis in supernatants of **(A)** *E. faecium* W54 and **(B)** *E. coli* BW25113 after adjusting the pH to different values. MPH hydrolysis in bacterial supernatants at different pH values, shown as the ratio of [RA]_24h_/[MPH]_0h_ quantified in ng/μL and normalized to d10-RA after 24 h of incubation with 50 μM MPH.

## Discussion

Although our initial experiments using small intestinal luminal content from rats **(Fig 1B)** suggested the capability of gut microbiota to metabolize MPH, our further observations of the spontaneous hydrolysis of MPH under physiological conditions (37 °C, pH 7.0)(**Fig 2B**), as well as the results from incubation of MPH in growth media in the presence and absence of pure bacterial cultures, uncovered the pH-dependent MPH hydrolysis, irrespective of the presence of bacteria **(Figs 3 and 4)**. The complex bacterial community present in the luminal content could have caused fluctuations in the pH levels during the 24 h period of incubation with MPH. Thus, we anticipate that the MPH hydrolysis observed when incubated with the small intestinal luminal content was mainly caused by an elevation of pH values during the duration of incubation.

To our knowledge, this is the first report that describes the effect of pH on the stability of MPH. Besides the analytical profile of MPH, where significant basic degradation was observed only at extreme temperatures (100 °C), [26], the stability of MPH at different pH levels has not been investigated. Significant MPH hydrolysis was shown at room temperature within only 30 min in a sodium hydroxide solution at extreme pH values [27][28]. Moreover, hydrolysis of MPH was also reported in static water [29], where MPH was completely hydrolyzed to RA within 37 h at 20 °C [29].

Changes in the pH along the gastrointestinal tract could induce non-enzymatic degradation of the MPH and therefore account, at least in part, for the low bioavailability of the drug. In a fasting state, MPH is predominantly absorbed in the jejunum, while under fed conditions absorption takes place mostly in the ileum [12]. Intraluminal pH changes along the gastrointestinal tract; the very acidic environment of the stomach rapidly changes in the SI, where pH increases up to 6 in the proximal SI and reaches 7.5 in the ileum [30]. This rise in the SI pH would result in 60% of MPH being hydrolyzed when it reaches the ileum **(Fig 3A)**. Preventing MPH from reaching an increased pH in the ileum by taking the medication under fasting conditions could improve its bioavailability, as MPH would be absorbed higher up in the small intestine where a pH below 7 should limit the non-enzymatic hydrolysis to around 10% **(Fig 3A)**. Reports on the pharmacokinetics of MPH comparing fed and fasting conditions are scarce and reveal contradictory results [31,32]. Although food intake tends to increase luminal pH of the gut [33], not enough information is available regarding the pH levels in the small intestine in fed or fasting conditions and how this could affect the bioavailability of MPH.

Microbial composition is another key factor that can influence the small intestinal pH and cause interindividual differences in MPH bioavailability. The gut microbiota is driven by the metabolism of dietary components[34]. For example, pH measured in ileostomy effluent from an ileostomy patient raised from 5.6 in the morning to 6.8 in the afternoon due to changes in feeding cycles [34] indicating that pH changes can indeed take place in the SI due to bacterial metabolism. Moreover, protein and amino acid deamination by gut bacterial metabolism results in the production of amine groups and ammonia that can also increase luminal pH [35]. Thus, interindividual differences in small intestinal microbial composition could be a key factor in MPH presystemic hydrolysis by shifting luminal pH either towards acidic pH, providing stability for MPH, or alkaline pH which would prompt MPH hydrolysis.

MPH metabolism was tested among a wide variety of drugs for their possible degradation by the gut microbiota [36]. MPH was among the drugs that were metabolized the least; only less than 10 colonic strains (which were not specified in the study) were able to metabolize around 20% of MPH at pH ≤ 6 [36]. Given that the majority of MPH is absorbed before it reaches the colon and considering bioavailability is around 30% [12], colonic bacterial metabolism cannot explain the low bioavailability of MPH in the ADHD patients.

Collectively, the present study shows that MPH is subject to spontaneous hydrolysis in a pH-dependent manner. The pH values at which MPH was hydrolyzed resemble the pH of the jejunum and ileum respectively, which are the main sites of absorption of MPH when administered orally [6]. Thus, differences in the intraluminal pH of the gastrointestinal tract rather than gut bacterial metabolization of could explain the low bioavailability of the ADHD main treatment, MPH. Moreover, the study provides a significant addition to previous studies reporting on the low bioavailability and interindividual variation in the response to MPH [7,8]. The main limitation of the current study is the lack of clinical measurements in ADHD patients to assess whether the interindividual variation in the response to MPH is related to differences among subjects in their small intestinal intraluminal pH caused by differences in diet, gut microbiota composition and their metabolic products, administration of other medications including antacids, MPH formulation administered, and whether the drug is taken in fasting or fed conditions.

## Acknowledgments

We thank Dr. Saskia van Hemert of Winclove Probiotics, Amsterdam, The Netherlands, for providing us *L. salivarius W1, L. plantarum W24 and E. faecium W54* strains, MEDICE Arzneimittel Pütter GmbH & Co KG. for financially contributing to the present research.

## Author Contributions

J.A., and S.E.A conceptualized and designed the study. J.A., W.M., and H.P., performed the experiments. J.A., H.P. and S.E.A. analyzed the data. J.A. and S.E.A. wrote the original manuscript that was reviewed by R.P, and J.P. Funding was acquired by S.E.A.

## Supplementary figures

**S1 Figure.**
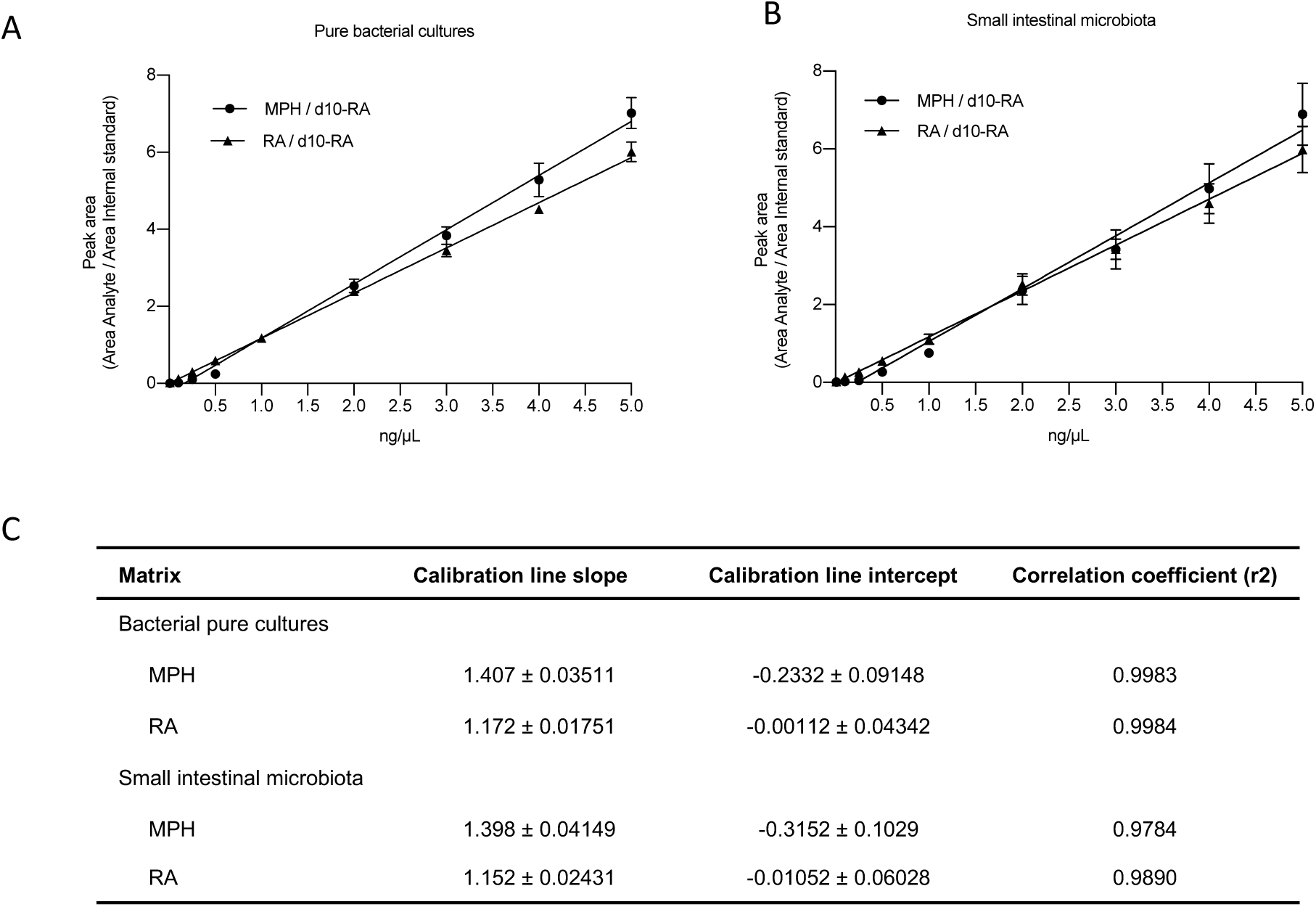
**(A, B)** Calibration curves obtained in the two different biological matrices used in this study: **(A)** pure bacterial cultures of *E. coli* BW25113 and **(B)** pool of small intestinal content of 5 WTG rats. Peak areas of methylphenidate (MPH) and ritalinic acid (RA) are normalized to the peak area of the internal standard d10-Ritalinic acid (d10-RA). **(C)** Linearity of the calibration curves fitted with a linear regression model. Data represents 3 biological replicates and error bars represent standard deviation.

**S2 Figure.**
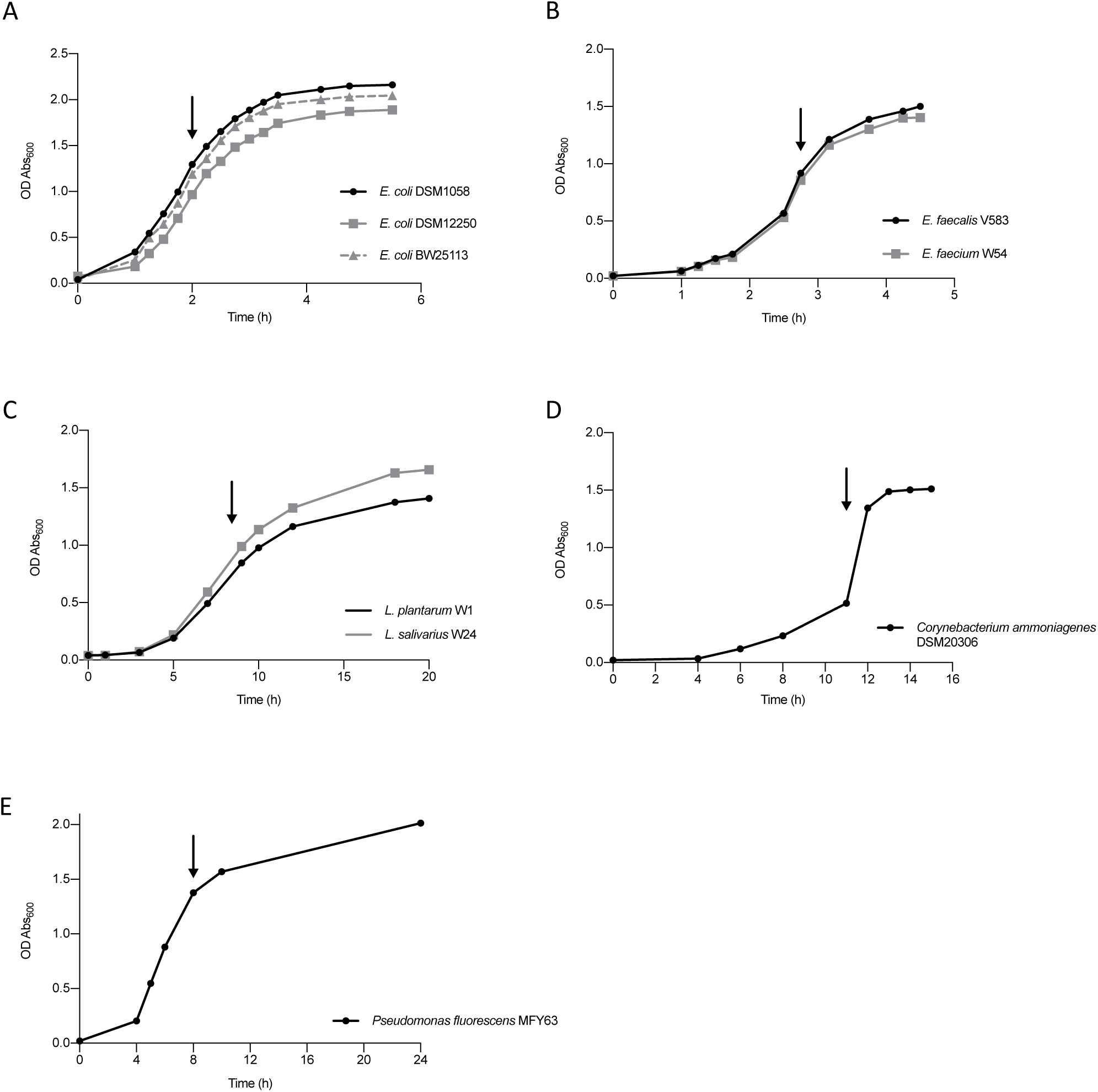
Growth curves of the strains used in this study. Optical density measured as the absorbance at 600 nm is plotted over time in aerobic cultures of **(A)** *E. coli* strains (BW25113, DSM11250 and DSM1058) grown at 37 °C, 220 rpm; **(B)** *Enterococcus* strains (*E. faecalis* V583 and *E. faecium* W54) strains grown at 37 °C without agitation **(C)** *Lactobacillus* strains (*L. plantarum* W1 and *L. salivarius* W24) grown at 37 °C, 220 rpm *Enterococcus* strains (*E. faecalis* V583 and *E. faecium* W54) strains grown at 37 °C without agitation; **(D)** *C. ammoniagenes* DSM20306 grown at 37 °C, 220 rpm and **(E)** *P. fluorescens* MFY63 grown at 37 °C, 220 rpm. Arrows indicate the late exponential phase when MPH was added.

**S3 Table.**
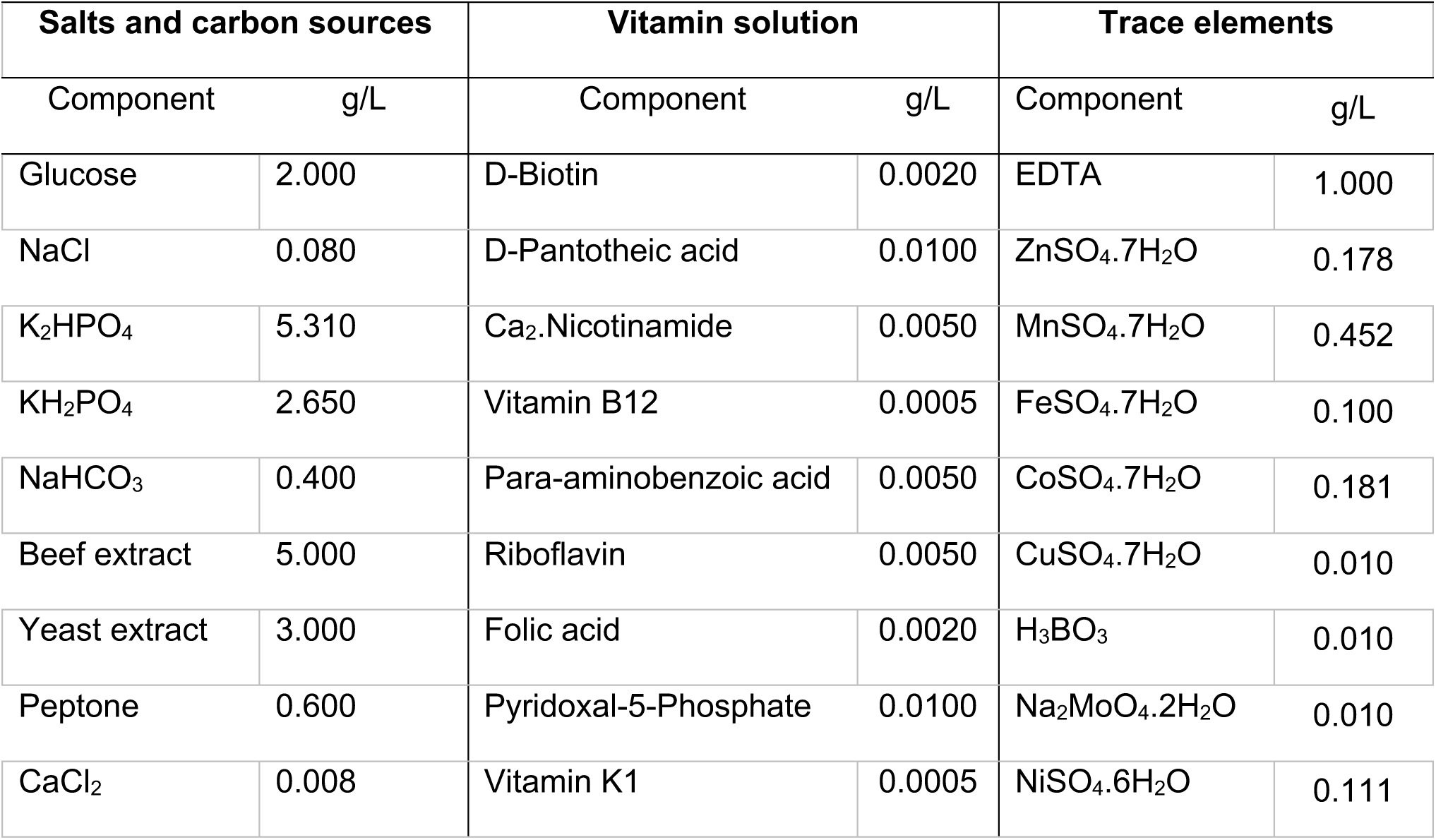
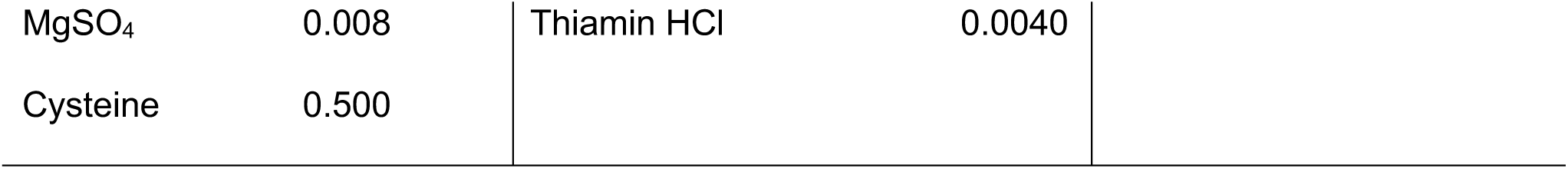
Constituents of enriched beef broth medium used in this study.

| Salts and carbon sources |  | Vitamin solution |  | Trace elements |  |
| --- | --- | --- | --- | --- | --- |
| Component | g/L | Component | g/L | Component | g/L |
| Glucose | 2.000 | D-Biotin | 0.0020 | EDTA | 1.000 |
| NaCl | 0.080 | D-Pantotheic acid | 0.0100 | ZnSO<sub>4</sub>.7H<sub>2</sub>O | 0.178 |
| K<sub>2</sub>HPO<sub>4</sub> | 5.310 | Ca<sub>2</sub>.Nicotinamide | 0.0050 | MnSO<sub>4</sub>.7H<sub>2</sub>O | 0.452 |
| KH<sub>2</sub>PO<sub>4</sub> | 2.650 | Vitamin B12 | 0.0005 | FeSO<sub>4</sub>.7H<sub>2</sub>O | 0.100 |
| NaHCO<sub>3</sub> | 0.400 | Para-aminobenzoic acid | 0.0050 | CoSO<sub>4</sub>.7H<sub>2</sub>O | 0.181 |
| Beef extract | 5.000 | Riboflavin | 0.0050 | CuSO<sub>4</sub>.7H<sub>2</sub>O | 0.010 |
| Yeast extract | 3.000 | Folic acid | 0.0020 | H<sub>3</sub>BO<sub>3</sub> | 0.010 |
| Peptone | 0.600 | Pyridoxal-5-Phosphate | 0.0100 | Na<sub>2</sub>MoO<sub>4</sub>.2H<sub>2</sub>O | 0.010 |
| CaCl<sub>2</sub> | 0.008 | Vitamin K1 | 0.0005 | NiSO<sub>4</sub>.6H<sub>2</sub>O | 0.111 |

|  |  |  |  |
| --- | --- | --- | --- |
| MgSO<sub>4</sub> | 0.008 | Thiamin HCl | 0.0040 |
| Cysteine | 0.500 |  |  |

